# High-quality genome assembly of a white eared pheasant individual and related functional and genetics data resources

**DOI:** 10.1101/2023.11.09.566452

**Authors:** Siwen Wu, Kun Wang, Tengfei Dou, Sisi Yuan, Dong-Dong Wu, Zhengchang Su, Changrong Ge, Junjing Jia

## Abstract

White eared pheasant (WT), (*Crossoptilon crossoptilon*), inhibiting at high altitudes (3000∼4,300 m), is a Galliformes bird native to the Qinghai, Sichuan, Yunnan and Tibet Province of China. Due to the difficulty of sequencing the precious species, there is no high-quality genome assembly for the species, hampering the understanding of their genetic mechanisms. To fill the gap, we sequenced and assembled a WT individual using Illumina short reads, PacBio long reads and Hi-C reads. With a contig N50 of 19.63 Mb, scaffold N50 of 29.59 Mb, total length of 1.02 Gb and BUSCO completeness of 97.2%, the assembly is highly complete. Evaluation shows that the assembly is at chromosome-level with only six gaps. Thus, our assembly provides a valuable genetic resource for the *crossoptilon* species. To further provide resources for gene annotation and population genetics analysis, we also sequenced transcriptomes of 20 tissues of the WT individual and re-sequenced another 10 individuals of WT. Our assembled WT genome and the sequencing data can be valuable resources to study the *crossoptilon* species.

## Background & Summary

*Crossoptilon*, belonging to the Phasianidae family in the Galliformes order, is a rare but important genus endemic in China [1]. There are four species belonging to the *Crossoptilon*, including Tibetan eared pheasant (TB) (C. harmani), brown eared pheasant (BR) (C. *mantchuricum*), white eared pheasant (WT) (C. *crossoptilon*) and blue eared pheasant (BL) (C. *auritum*) [2, 3]. TBs are only distributed in southeastern Tibet with high altitude (more than 6,000 m), BRs are mainly distributed in mountains of Beijing, Shanxi and Hebei provinces with low altitude (20∼1,000 m) [2], BLs are only encountered in the mountains of Qinghai, Gansu and Sichuan provinces and Ningxia Autonomous Region with normal altitude (1,500∼3,000 m) [2], and WTs are distributed in Qinghai, Sichuan, Yunnan and Tibet Province of China with relatively high altitude (3,000∼7,000 m) [2]. All the four species are national key protection animals in China and are of high commercial value. However, studies related to the four species are rare, and most are limited to the study of single genes, partial sequences [4, 5] or mitochondrial DNA sequences [1, 6]. Due to the bottleneck period of *Crossoptilon* in history, only studying maternal genetic information could lead to inaccurate results [7]. Therefore, it is more important to study the phylogeny and taxonomy of *Crossoptilon* species using the whole genome sequences. The genome of a BR individual was assembled in 2020 [8], however, since they only used the Illumina short reads and fragment libraries to assemble the genome, the contig N50 and scaffold N50 is only 0.11 Mb and 3.63 Mb, respectively, and the BUSCO complete value [9] is only 95.1%, which was not continuous and accurate enough to study the *Crossoptilon* species.

Thus, to elucidate genomic features of these vulnerable species, we assembled the genome of a WT individual using the Illumina short reads, PacBio long reads and Hi-C reads. The resulting assembly has a total length of 1.02 Gb, with a contig N50 of 19.6 Mb, a scaffold N50 of 29.6 Mb, a complete BUSCO value [9] of 97.2% and only six gaps. Other aspects of evaluation also indicate that our assembly is at chromosome-level with very high quality. Thus, our assembly provides a good reference assembly for the *Crossoptilon* species and would help the conservation planning for the threatened species. Moreover, we also sequenced the transcriptomes of 20 tissues of the WT individual and re-sequenced another 10 individuals of WT, which can be valuable resources for studying the biology, evolution and developing conservation strategies of these endangered species.

## Methods

### Bird populations

For the WT populations, blood samples of a total of 11 WT individuals (five males and six females) aged 10 months were collected from Diqing Tibet Autonomous prefecture, Yannan, China. A female individual was collected from the same area for whole genome assembly and its relevant tissues were subject to Illumina paired-end DNA short reads sequencing, PacBio long reads sequencing, Hi-C paired-end short reads sequencing, and paired-end RNA-seq of 20 tissues (Heart, Liver, Spleen, Lung, Kidney, Pancreas, Gizzard, Glandular, Crops, Ovary, Abdominal fat, Rectum, Duodenum, Cecum, Skin, Small intestine, Brain, Cerebellum, Chest muscle, Leg muscle) and mixed tissues for gene annotation. All other 10 individuals were subject to Illumina paired-end DNA short reads sequencing.

### Ethics approval

All the experimental procedures were approved by the Animal Care and Use Committee of the Yunnan Agricultural University (approval ID: YAU202103047). The care and use of animals fully complied with local animal welfare laws, guidelines, and policies.

### Short reads DNA sequencing

Two milliliters of blood were drawn from the wing vein of each bird in a centrifuge tube containing anticoagulant (EDTA-2K) and stored at -80°C until use. Genomic DNA (10µg) in each blood sample was extracted using a DNA extraction kit (DP326, TIANGEN Biotech, Beijing, China) and fragmented using a Bioruptor Pico System (Diagenode, Belgium). DNA fragments around 350 bp were selected using SPRI beads (Beckman Coulter, IN, USA). DNA-sequencing libraries were prepared using Illumina TruSeq® DNA Library Prep Kits (Illumina, CA, USA) following the vendor’s instructions. The libraries were subject to 150 cycles paired-end sequencing on an Illumina Novaseq 6000 platform (Illumina, CA, USA) at 102X coverage.

### PacBio long reads DNA sequencing

High molecular weight DNA was extracted from each blood sample using NANOBIND® DNA Extraction Kits (PacBio, CA, USA) following the vendor’s instructions. DNA fragments of about 25 kb were size-selected using a BluePippin system (Sage Science, MA, USA). Sequencing libraries were prepared for the DNA fragments using SMRTbell® prep kits (PacBio, CA, USA) following the vendor’s instructions, and subsequently sequenced on a PacBio Sequel II platform (PacBio, CA, USA) at 91X coverage.

### Transcriptome sequencing

One to two grams of various tissues and mixed tissues were collected from the selected female WT individual in a centrifuge tube and immediately frozen in liquid nitrogen, then stored at -80°C until use. Total RNA from each tissue sample were extracted from each tissue or mixed tissues using TRlzol reagents (TIANGEN Biotech, Beijing China) according to the manufacturer’s instructions. RNA-sequencing libraries for each tissue collected from the individual were prepared using Illumina TruSeq® RNA Library Prep Kits (Illumina, San Diego) following the vendor’s instructions. The libraries were subject to 150 cycles paired-end sequencing on an Illumina Novaseq 6000 platform at a sequencing depth of 281X.

### Hi-C reads sequencing

Five milliliters of blood were drawn from the wing vein of the selected WT individual in a Streck Cell-free DNA BCT collecting vessel (Streck Corporate, USA), and stored at 4°C and used in 24 hours. Hi-C libraries were constructed using Phase Genomics’ Animal Hi-C kit following the vendor’s instructions and subsequently sequenced on an Illumina’s Novaseq 6000 platform at a sequencing depth of 81X.

### Quality assessment of sequencing data

We used FastQC (0.12.1) [10] to evaluate the quality of the Illumina DNA short reads, PacBio long reads, Hi-C reads, RNA-seq reads and re-sequencing reads of the WT.

### Contig assembling and scaffolding

We used the PacBio long reads longer than 5,000 bp to assemble the contigs using Wtdbg (2.5) [11], and polished the contigs using Wtdbg (2.5) [11] with Illumina DNA short reads for the WT. Then we used SALSA [12, 13] to bridge the contigs and obtain the scaffolds with Hi-C short reads. We filled the gaps in the scaffolds using PBJelly [14] with the PacBio long reads, and then made two rounds of polish by firstly using Racon (1.4.21) [15] with PacBio long reads and secondly using NextPolish (1.4.0) [16] with Illumina DNA short reads from the selected WT individual.

### Quality evaluation of assemblies

We masked the repeats for the assembly of the WT using WindowMasker (2.11.0) [17] to get the repeat rate, and estimated the heterozygosity of the assembly using Jellyfish (2.3.0) [18] and GenomeScope [19]. To estimate the continuity of the assembly, we used QUAST (5.0.2) [20] to calculate the contig N50 and scaffold N50. To estimate the structural accuracy, we used Asset [21] to calculate the reliable block N50 and used BUSCO (5.1.3) [9] to calculate the false duplication rate for the assembly. To estimate the base accuracy, we used Merqury (1.3) [22] to calculate the k-mer QV and k-mer completeness for the assembly, used BWA (0.7.17) [23] to map the short reads of the selected WT individual to the assembly, and used SAMtools (1.10) [24] to analyze the mapping results. To estimate the functional completeness, we used BUSCO (5.1.3) [9] to assess the completeness of the assembly against the avian gene set. To plot the heatmap of the scaffolds of the assembly, we mapped the Hi-C paired-end short reads to the assembly using BWA (0.7.17) [23], used SAMtools (1.10) [24] and Pairtools (0.3.0) [25] to analyze the mapping results, and used Higlass [26] to plot the heatmap for the assembly.

### Data Records

The Illumina DNA paired-end short reads, PacBio long reads, Hi-C paired-end short reads and the RNA-seq paired-end short reads of different tissues of the selected WT individual are available at NCBI SRA with the accession number PRJNA956489 (https://www.ncbi.nlm.nih.gov/bioproject/?term=PRJNA956489), and the genome assembly of the individual is available at FigShare (https://figshare.com/articles/dataset/High-quality_genome_assembly_of_a_white_eared_pheasant_individual/24535015). The re-sequencing paired-end short reads of the other 10 white eared pheasant individuals are available at NCBI SRA with the accession number PRJNA956570 (https://www.ncbi.nlm.nih.gov/bioproject/?term=PRJNA956570).

### Technical Validation

#### Quality evaluation of the sequencing data of the WT

We generated Illumina DNA paired-end short reads, PacBio long reads and Hi-C paired-end short reads for a female WT individual. As shown in Table 1, for the Illumina DNA paired-end short reads, the sequencing length is 150 bp and the sequencing depth is 102X. For the PacBio long reads, the average sequencing length is 10 kbp and the sequencing depth is 91X. For the Hi-C paired-end short reads, the sequencing length is 150 bp and the sequencing depth is 81X. In addition, we also sequenced the transcriptomes of 20 tissues of the WT individual and re-sequenced another 10 individuals of WT. As shown in Table 1, for the RNA-seq reads, the sequencing length is 150 bp and the sequencing depth is 281X. For the re-sequencing reads, the sequencing length is 150 bp and the average sequencing depth is 62X.

**Table 1:** Raw sequencing data of the WT individual.

|  |  |  |
| --- | --- | --- |
| **Short reads** | **Depth** | 102X |
|  | **Length** | 150 bp |
|  | **# Pairs** | 338,515,576 |
| **Long reads** | **Depth** | 91X |
|  | **Average reads length** | 10 kbp |
|  | **# Total reads** | 8,982,129 |
|  | **# Reads > 5 kbp** | 5,557,754 |
| **Hi-C reads** | **Depth** | 81X |
|  | **Length** | 150 bp |
|  | **# Pairs** | 269,697,038 |
| **RNA-seq reads** | **Depth** | 281X |
|  | **Length** | 150 bp |
|  | **# Tissues** | 20 |
|  | **# Pairs** | 936,233,391 |
| **Re-sequencing** | **# Individuals** | 10 |
|  | **Average depth** | 62X |
|  | **Length** | 150 bp |

Figures 1a∼1e show the quality assessment of the different sequencing data of WT. All the Illumina DNA paired-end short reads, Hi-C paired-end reads, RNA-seq reads and re-sequencing reads have a Phred score greater than 35 (Figures 1a∼1d), suggesting that the base accuracy of all these reads is greater than 99.9% [27] and are of very high quality. For the PacBio long reads, since they do not come with Phred quality scores, the quality was evaluated using length distribution (Figure 1e), which indicates the average length is about 10 kbp and is of high quality.

**Figure 1.**
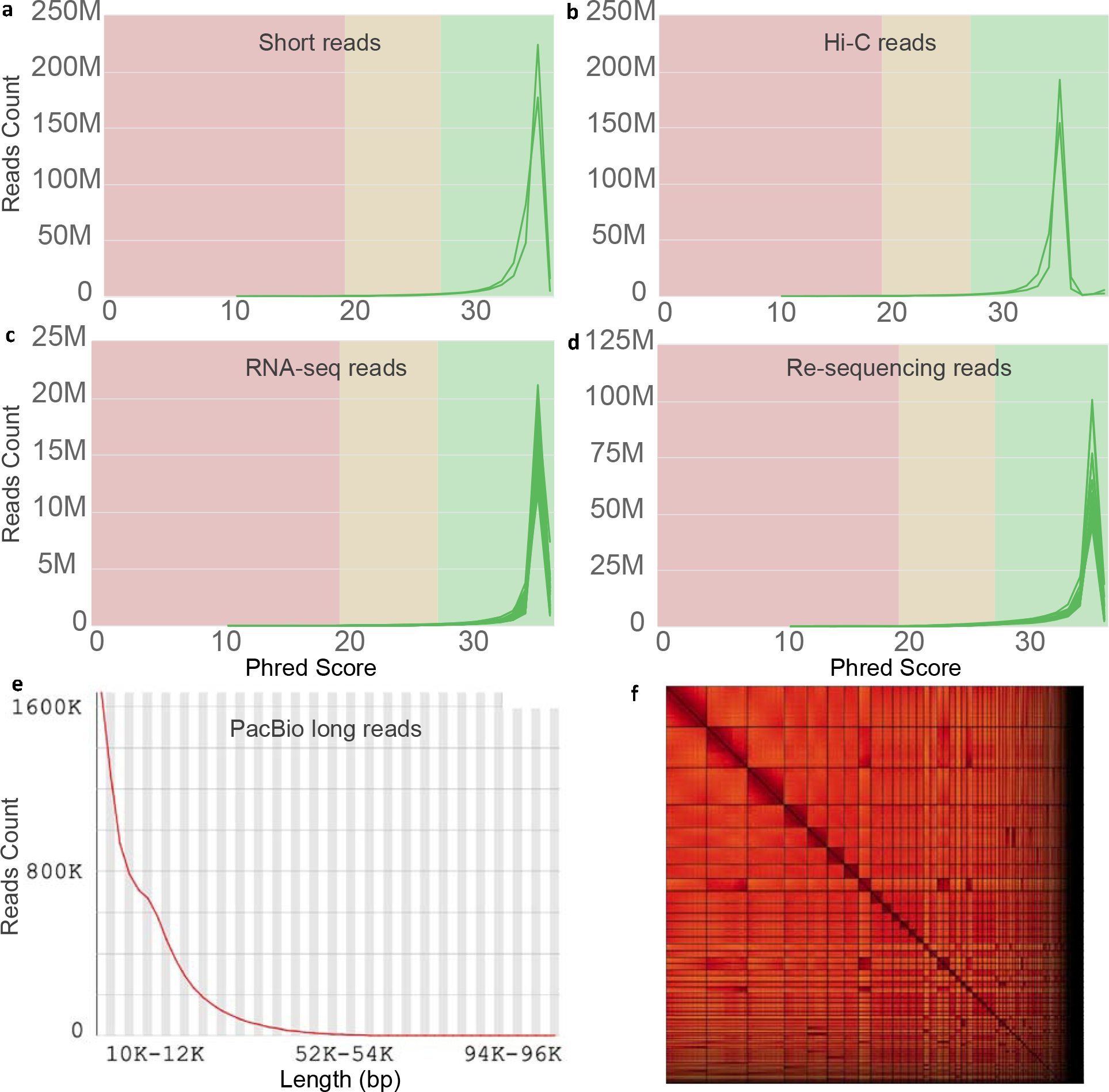
Quality assessment of different types of sequencing reads of the WT and Hi-C interaction heatmap of the scaffolds of the WT assembly.

### Evaluation of the assembly quality of the WT

Using the short and long sequencing reads, we assembled the genome of a WT individual into 805 contigs with a contig N50 of 19.63 Mb. The total length of the contigs is 1.02 Gb, comparable to those of the chicken (*Gallus gallus*) genome assemblies GRCg6a and GRCg7b/w as well as of the previously assembled BR genome [8] (1.01Gb) (Table 2). Using the Hi-C paired-end short reads, we further assembled the contigs into 643 scaffolds with a scaffold N50 of 29.59 Mb (Table 2). We assessed the quality of the assembly using the criteria proposed by the VGP consortium [28], and compared it with chicken assemblies GRCg6a and GRCg7b/w, the best-studied bird genomes. These criteria include genome features (heterozygosity and repeat rates), continuity (assembly size, N50 and gaps), structure accuracy (reliable block N50 and false duplication rate), base accuracy (k-mer QV, k-mer completeness and short reads mapping rate) and functional completeness (BUSCO completeness) (Table 2).

**Table 2:** Evaluation of the genome assembly of the WT individual.

| Breed | Genome |  | Continuity |  |  |  | Structural Acc. |  | Base Acc. |  |  | Func. Comp. |
| --- | --- | --- | --- | --- | --- | --- | --- | --- | --- | --- | --- | --- |
|  | Het (%) | Rep (%) | Size (Gb) | # Contigs (Contig N50) (Mb) | # Scaffolds (Scaffold N50) (Mb) | # Gaps | Reliable block N50 (Mb) | False duplications (%) | k-mer QV | k-mer Comp. (%) | Short reads Comp. (%) | BUSCO Comp. (%) |
| WT | 0.54 | 20.6 | 1.023 | 805 (19.6) | 643 (29.6) | 6 | 14.6 | 0.3 | 42.0 | 95.3 | 99.3 | 97.2 |
| GRCg6a | - | 20.6 | 1.056 | 1,402 (17.7) | 464 (20.8) | 500,945 | - | 0.4 | - | - | - | 96.6 |
| GRCg7b | - | 20.6 | 1.050 | 677 (18.8) | 214 (90.9) | 463 | - | 0.4 | - | - | - | 96.6 |
| GRCg7w | - | 20.2 | 1.046 | 685 (17.7) | 276 (90.6) | 409 | - | 0.4 | - | - | - | 96.8 |

The heterozygosity rate of the WT is 0.54%, and its repeat rate is 20.6%, both are comparable to those of the GRCg6a and GRCg7b/w assemblies (Table 2). For the continuity, the contig N50 (19.6 Mb) of the assembly is slightly larger than those of the GRCg6a and GRCg7b/w assemblies. The scaffold N50 (29.6 Mb) of the assembly is slightly larger than that of the GRCg6a assembly, but smaller than those of the GRCg7b/w assemblies (Table 2). For gaps, there are only six gaps in our assembly, which is much fewer than those of the GRCg6a and GRCg7b/w assemblies (Table 2), indicating that our assembly is almost gapless. For the structural accuracy, the reliable block N50 [21] of our assembly (14.6 Mb) is comparable to those of Avian genomes assembled by the recent VGP consortium [28]. The false duplication rate [9] of our assembly (0.3%) is slightly smaller than those of the GRCg6a and GRCg7b/w assemblies (Table 2), indicating that the structural accuracy of our assembly is very high. For the base accuracy, the k-mer QV of our assembly is 42.0, suggesting that the consensus base accuracy is greater than 99.99% [22] (Table 2). The k-mer completeness (defined as the fraction of reliable k-mers in highly accurate short reads data that are also found in the assembly [22]) of our assembly is 95.3% (Table 2), which is comparable to those of the recent VGP assemblies [28]. To further evaluate the base accuracy, we mapped the Illumina short reads of the WT individual to the assembly and found that 99.3% short reads can be mapped to the assembly (Table 2), suggesting that our assembly is of high base accuracy. For the functional completeness, we achieved a larger BUSCO completeness value [9] (97.2%) than those of the GRCg6a and GRCg7b/w assemblies (Table 2), suggesting that our assembly is of high functional completeness. To further check whether our assembly is at chromosome-level, we plotted the Hi-C interaction heatmap of the scaffolds. As shown in Figure 1f, almost all the scaffolds form a square at the diagonal of the heatmap, indicating that our assembly is at chromosome-level, although we lack genetic marks to sort them into specific chromosomes.

## Code Availability

All genome assembly code and the corresponding pipeline description are available at https://github.com/zhengchangsulab/A-genome-assebmly-and-annotation-pipeline.

## Author Contributions

ZS, JJ, CG and DW supervised and conceived the project; KW and TD collected tissue samples and conducted molecular biology experiments; SW and SY^1^ assembled the genomes; SW and ZS performed data analysis; and SW, KW, ZS and JJ wrote the manuscript.

## Acknowledgments

This work was supported by the National Natural Science Foundation of China (U2002205 and U1702232), Yunling Scholar Training Program of Yunnan Province (2014NO48), Yunling Industry and Technology Leading Talent Training Program of Yunnan Province (YNWR-CYJS-2015-027), Natural Science Foundation of Yunnan Province (2019IC008 and 2016ZA008), and Department of Bioinformatics and Genomics of the University of North Carolina at Charlotte.

